# FasR regulates fatty acid biosynthesis and is essential for virulence of *Mycobacterium tuberculosis*

**DOI:** 10.1101/2020.06.08.140004

**Authors:** Sonia Mondino, Cristina L. Vázquez, Matías Cabruja, Claudia Sala, Amaury Cazenave-Gassiot, Federico C. Blanco, Markus R. Wenk, Fabiana Bigi, Stewart T. Cole, Hugo Gramajo, Gabriela Gago

**Affiliations:** Laboratory of Physiology and Genetics of Actinomycetes, Instituto de Biología Molecular y Celular de Rosario (IBR-CONICET), Facultad de Ciencias Bioquímicas y Farmacéuticas, Universidad Nacional de Rosario, Rosario, Argentina; Instituto de Biotecnología-IABIMO, UEDD CONICET-Instituto Nacional de Tecnología Agropecuaria (INTA), Hurlingham, Argentina; Global Health Institute, Ecole Polytechnique Fédérale de Lausanne, Lausanne, Switzerland; Singapore Lipidomics Incubator (SLING), Life Sciences Institute, and Department of Biochemistry, Yong Loo Lin School of Medicine, National University of Singapore, Singapore

**Author notes:** Correspondence, Gabriela Gago. Institut Pasteur, Paris (France) (Sonia Mondino, Stewart T. Cole). Stanford University (United States) (Matías Cabruja). Fondazione Toscana Life Sciences, Siena (Italy) (Claudia Sala). Sonia Mondino, Cristina L. Vázquez and Matías Cabruja contributed equally to this work.

**Keywords:** tuberculosis, fatty acid biosynthesis, lipid homeostasis, mycobacterial virulence, mycobacterial cell envelope

## Abstract

*Mycobacterium tuberculosis*, the etiologic agent of human tuberculosis, is the world’s leading cause of death from an infectious disease. One of the main features of this pathogen is the complex and dynamic lipid composition of the cell envelope, which adapts to the variable host environment and defines the fate of infection by actively interacting with and modulating immune responses. However, while much has been learned about the enzymes of the numerous lipid pathways, little knowledge is available regarding the proteins and metabolic signals regulating lipid metabolism during *M. tuberculosis* infection. In this work, we constructed and characterized a FasR-deficient mutant in *M. tuberculosis* and demonstrated that FasR positively regulates *fas* and *acpS* expression. Lipidomic analysis of the wild type and mutant strains revealed complete rearrangement of most lipid components of the cell envelope, with phospholipids, mycolic acids, sulfolipids and phthiocerol dimycocerosates relative abundance severely altered. As a consequence, replication of the mutant strain was impaired in macrophages leading to reduced virulence in a mouse model of infection. Moreover, we show that the *fasR* mutant resides in acidified cellular compartments, suggesting that the lipid perturbation caused by the mutation prevented *M. tuberculosis* inhibition of phagolysosome maturation. This study identified FasR as a novel factor involved in regulation of mycobacterial virulence and provides evidence for the essential role that modulation of lipid homeostasis plays in the outcome of *M. tuberculosis* infection.

## Introduction

Tuberculosis (TB) is one of the top ten causes of death and the leading cause from a single infectious agent, exceeding both malaria and HIV. In 2018, there were an estimated 10 million new cases of active TB and 1.3 million patients died from active infection (WHO, 2019). The emergence of multidrug-resistant TB (MDR-TB) presents an increasingly difficult therapeutic challenge; consequently, MDR-TB is now the main cause of death due to antimicrobial resistance (WHO, 2019) thus transforming this long neglected disease into a global health priority.

The composition and complexity of the cell envelope is the most distinctive feature of *Mycobacterium tuberculosis*, the etiologic agent of human TB. Biochemical and microscopic data revealed that the mycobacterial envelope consists of three discrete entities: a plasma membrane surrounded by a complex cell wall and an outermost layer called the capsule in pathogenic species (Chiaradia et al., 2017; Daffé and Marrakchi, 2019). The *M. tuberculosis* cell wall consists of peptidoglycan covalently linked to arabinogalactan esterified by mycolic acids (MA), which form the inner leaflet of the outer membrane bilayer (mycomembrane). The outer leaflet of this membrane is composed of free, non-covalently bound lipids and glycolipids like trehalose monomycolates (TMM), trehalose dimycolates (TDM), glycerol monomycolates, glucose monomycolates, phthiocerol dimycocerosates (PDIM), poly acylated trehaloses (PAT), sulfolipids (SL), phosphatidylinositol mannosides (PIM), phenolic glycolipids (PGL) and mannose-capped lipoarabinomannans (Man-Lam) (Brennan and Nikaido, 1995; Daffe and Reyrat, 2008).

During the course of infection, intracellular pathogenic bacteria use different strategies to survive, multiply and persist inside the host (Russell et al., 2010). *M. tuberculosis* encounters heterogeneous host environments, including high oxygen tension in the lung alveolus, reactive oxygen and nitrogen species in activated macrophages, as well as nutrient depletion and hypoxia in tuberculous granulomas. How *M. tuberculosis* survives these diverse environments is not fully understood. *M. tuberculosis* has evolved a wide array of specific lipids and related metabolic pathways that actively interact with the immune system, and multiple reports from human and animal studies suggest an intimate link between *M. tuberculosis* lipid metabolism and outcome of infection (Neyrolles and Guilhot, 2011; Passemar et al., 2014; Vromman and Subtil, 2014). Of particular interest is the challenge that *M. tuberculosis* must face in order to adjust its own lipid metabolism while the macrophage is changing its lipid content (Teng et al., 2017). Little is known about the mechanism underlying the reorganization of *M. tuberculosis* envelope during infection (Dulberger et al., 2020). Understanding how this pathogen selects and organizes the envelope components to resist killing, and how these components make the envelope essential to *M. tuberculosis* survival, virulence and pathogenesis, is still a major challenge. Therefore, characterization of the regulatory elements that control these processes should help improve understanding of the mechanisms involved in mycobacterial pathogenicity and could provide new potential targets for anti-TB chemotherapy.

Previous studies on the regulation of lipid biosynthesis in mycobacteria allowed us to identify FasR as the transcriptional activator of fatty acid biosynthesis in *M. smegmatis* (Mondino et al., 2013). Deletion of *fasR* from *M. smegmatis* was possible only in the presence of an extra copy of the gene, suggesting that *fasR* is essential for viability. Notably, we found that the affinity of FasR for its target promoter is modulated by long chain acyl-CoAs (≥C_16_), the end products of the FAS I pathway. FasR, together with MabR, the transcriptional regulator of MA biosynthesis, are the only transcription factors involved in fatty acid biosynthesis regulation known to be essential for *M. smegmatis* survival, reflecting the relevance of lipid homeostasis for mycobacterial viability (Salzman et al., 2010; Mondino et al., 2013).

In this study, a *M. tuberculosis* knockout strain in *fasR* was constructed and the mutant used to analyze the implications of lipid homeostasis in *M. tuberculosis* virulence. Inactivation of *fasR* in *M. tuberculosis* was not lethal for *in vitro* growth but led to complete rearrangement of lipid composition of the envelope. Consequently, the mutant strain was less virulent in a mouse model and showed impaired replication in macrophages, thus implying that lipid homeostasis plays an important role during *M. tuberculosis* infection.

## Materials and methods

### Ethics statement

All animal experiments were performed in compliance with the regulations of the Institutional Animal Care and Use Committee (CICUAE) of INTA with the authorization Number 20/2016.

### Bacterial strains, culture and transformation conditions

The *E. coli* strain DH5α (Hanahan, 1983) was used for routine subcloning and was transformed according to Sambrook *et al*. (Sambrook, 1989). *M. tuberculosis* H37Rv (Cole et al., 1998), *Mtb*Δ*fasR* and *Mtb*Δ*fasR*-c*fasR* were grown at 37 °C in Middlebrook 7H9 broth (DIFCO) supplemented with 0.2 % glycerol, 0.03 % Tyloxapol and 10 % OADC or Middlebrook 7H10 agar (DIFCO) supplemented with 0.2 % glycerol and 10 % OADC. Hygromycin (Hyg, 50 μg/ml), kanamycin (Kan, 15 μg/ml) and streptomycin (Str, 12 μg/ml) were added when needed.

Recombinant plasmids and strain genotypes are listed in electronic supplementary material (Tables S1 and S2).

### DNA manipulation, plasmid construction and mutant generation

Isolation of plasmid DNA, restriction enzyme digestion and agarose gel electrophoresis were carried out by conventional methods (Sambrook, 1989). Genomic DNA of *M. tuberculosis* was obtained as described previously (Connell, 1994). For plasmid generation purposes, all PCR products were ligated into the pCR^®^-Blunt II-TOPO^®^ Vector (Invitrogen) according to the manufacturer’s recommendation.

For the construction of the *fasR* mutant allele, a fragment of 946 bp containing 58 bp of the *fasR* 5’end plus 882 bp of the upstream region was PCR amplified from genomic DNA using oligonucleotides 3208_1up and 3208_2down (Table S3). The PCR product was cloned in the pCR BluntII TOPO vector (Invitrogen). The resulting plasmid was digested first with *Eco*RV and after with *Xho*I/*Nde*I. The 940 bp fragment obtained was cloned in the suicide vector pJG1100 (Gomez and Bishai, 2000; Muñoz-Elías and McKinney, 2006) digested with the same enzymes, generating plasmid pFR34. Additionally, a fragment of 883 bp containing 87 bp of the *fasR* 3’end plus 784 bp of the downstream region was PCR amplified from genomic DNA using oligonucleotides 3208_3up and 3208_4down (Table S3). The PCR product was cloned in the pCR BluntII TOPO vector (Invitrogen). The resulting plasmid was digested first with *Eco*RI and after with *Nde*I/*Spe*I and the fragment (877 bp) was cloned in pFR34 vector digested with the same enzymes, generating plasmid pFR35.

For the construction of a *fasR* merodiploid strain, the fragment resulting from the digestion of the plasmid pFR3 (Mondino et al., 2013) with *Xba*I and *Eco*RI was cloned between the *Avr*II and *Mfe*I sites in plasmid pGA44 (Kolly et al., 2014). The resulting plasmid, pFR33, is an integrative plasmid that harbours *fasR* expressed under the control of *Ptr* promoter and the TetR/Pip OFF system necessary for its regulation.

Deletion of *fasR* was accomplished by homologous recombination. After transformation of *M. tuberculosis* with plasmid pFR35, the first recombination event was selected on 7H10 medium containing Kan and Hyg and resistant colonies were screened by colony PCR. The merodiploid strain was generated by integration of pFR33 at the L5 *attB* site and transformants were plated on 7H10 with Hyg, Kan and Str. Deletion of the wild type gene by allelic replacement was accomplished by plating bacteria on 7H10 agar plates supplemented with Str and 2.5 % sucrose. The Hyg^s^, Kan^s^, Str^r^, Sucrose^r^ colonies were tested by PCR and the *fasR* conditional knockdown strain was named *Mtb*Δ*fasR*-c*fasR*. Generation of a *fasR* knockout strain (*Mtb*Δ*fasR*) was obtained by swapping the pFR33 plasmid at the L5 *attB* site with the pND255 plasmid. Transformants were selected on 7H10 plates containing Hyg. Deletion of *fasR* was confirmed by Southern Blot and colony PCR.

### Southern Blot

10 μg of genomic *M. tuberculosis* DNA was digested overnight with an excess of *Xho*I, and the fragments were separated by electrophoresis in a 0.7% agarose gel. Southern blotting was carried out using Hybond-N^+^ membrane (Amersham) and DIG High Prime DNA Labeling and Detection Starter kit II (Roche), following the indications of the manufacturer. The probe consisted of a 594 bp fragment amplified with oligonucleotides Southern_1fasRTB and Southern_2fasRTB (Table S3), corresponding to the upstream genomic region of *fasR*.

### Western blot

Equal amounts of protein extracts were loaded on 4-12 % NuPAGE gels (Invitrogen) and transferred onto nitrocellulose membranes using a dry electrophoresis transfer apparatus (Invitrogen). Membranes were incubated in TBS-Tween blocking buffer (20 mM Tris pH 7.5, 150 mM NaCl, 0.1 % Tween 20) with 5 % (w/v) skimmed milk powder for 2 h at 4 °C prior to overnight incubation with 1:1000 dilution of monoclonal anti-FasR primary antibody (Alere™). Membranes were washed in TBS-Tween three times at room temperature, and then incubated with 1:100000 secondary antibody Anti-Mouse Kappa conjugated to Horseradish Peroxidase (Southern Biotech) for 2 h before washing again. Signals were detected using Chemiluminescent Peroxidase Substrate (Sigma-Aldrich). Monoclonal mouse anti-RpoB antibodies were purchased from NeoClone and used 1:20000.

### RNA techniques

RNA was extracted from *M. tuberculosis* strains grown for 8 days in complete 7H9 media using Trizol-phenol based methods (Kolly et al., 2014) and treated, when needed, with RQ1 RNase-Free DNase (Promega). qRT-PCR was performed using second strand cDNA as the template, generated with SuperScript III Reverse Transcriptase (Invitrogen), random primers and a green fluorochrome as the indicator dye (qPCR master mix, Biodynamics). The expression of *fas, fasR* and *acpS* was quantified after normalization of RNA levels to the expression of the *sigA* gene as previously described (Cabruja et al., 2017). qPCR data are presented as relative expression values normalized to *sigA* and to the wild type strain H37Rv. qPCR cycling conditions were as follows: 95 °C for 2 min followed by 40 cycles of 95 °C for 15 s, 58 °C for 15 s and 68 °C for 20 s. The sequences of all the primers used are listed in electronic supplementary material (Table S3).

### Lipid analysis

Cell-associated lipid for lipidomic analyses were prepared as previously described (Cabruja et al., 2017). Briefly, cell pellets of cultures grown at 37 °C in Middlebrook 7H9 broth (DIFCO) supplemented with 0.2 % glycerol, 0.03 % Tyloxapol and 10 % OADC were washed with 50 mM ammonium acetate, pH 7.8 and transferred to glass tubes containing 5 ml CHCl_3_:CH_3_OH (2:1, v/v). Samples were incubated overnight at 4 °C with gentle agitation. After centrifugation, bacterial pellets were subjected to an additional extraction using CHCl_3_:CH_3_OH (1:2, v/v) for 2 h. Organic extracts were pooled and dried under nitrogen at 4 °C. Lipids were resuspended in 3 ml of CHCl_3_, washed with 3 ml of H_2_O and the organic phase was transferred to pre-weighted glass tubes, dried under nitrogen at 4 °C and re-weighted on a microbalance. Extracts were dissolved in CHCl_3_:CH_3_OH (1:1, v/v) at 1 mg ml^-1^ and centrifuged at 3000 g for 5 min. Lipids were analyzed in an Agilent 1200 series HPLC system with a Reprospher (Dr Maish) 100 C8 column (1.8 μm, 50 mm length, 2 mm ID). The flow rate was 0.3 ml min^-1^ in binary gradient mode with the following elution program: the column was equilibrated with 100 % mobile phase A [CH_3_OH:H_2_O (99:1, v/v), containing 0.05 mM AcNH_4_]. 2 μl of each sample were injected, and the same elution conditions continued for 1 min, followed by a 12 min gradient to 100 % mobile phase B [isopropanol:hexane:H_2_O (79:20:1, v/v), containing 0.05 mM AcNH_4_], finally holding this condition for 1 min. The column was equilibrated for 2 min with 100 % mobile phase A before injection of the next sample. An Agilent 6500 series Q-TOF instrument with a Dual AJS ESI was used for mass analysis. Ionization gas temperature was maintained at 200 °C with a 14 l min^-1^ drying gas flow, a 35 psig nebulizer pressure and 3500 Volts. Spectra were collected in positive and negative mode from *m/z* 115 to 3000 at 4 spectra s^-1^. Automatic tandem-MS (MS-MS) was obtained in data-dependent acquisition mode at 15 MS-MS per cycle. Continuous infusion calibrants included *m/z* 121.051 and 922.010 in positive-ion mode and *m/z* 119.035 and 955.972 in negative-ion mode. CID-MS was carried out with an energy of 50 volts. For the comparative analysis, the column is conditioned by four successive mock injections before randomized QCs and mycobacterial samples are analyzed. Lipid features were annotated based on structural MS-MS information. Lipid classes’ retention times (RT) were identified using characteristics fragmentation patterns (Layre et al., 2011). Individual molecules in each class were then identified based on exact mass and RT.

### Colony-forming units (CFU) analysis

*M. tuberculosis* H37Rv, *Mtb*Δ*fasR* and *Mtb*Δ*fasR*-c*fasR* strains were used to infect THP-1 cells (MOI=5). The day before infection 4×10^5^ THP-1 cells were plated in a 24-well plate in RPMI Medium with 4.5 g/L glucose, 10 % heat inactivated fetal calf serum (Internegocios) and 2 mM L-glutamine (full medium) plus 50 nM phorbol 12-myristate 13-acetate (PMA) to allow differentiation of the cells to macrophages. On the infection day, cells were washed and a suspension of the different strains of *M. tuberculosis* in RPMI medium was added. After 2 h of uptake, cells were washed 3 times with PBS and fresh full RPMI medium plus gentamicin 10 μg mL^-1^ was added. Cells were incubated at 37 °C in 5 % CO_2_ atmosphere. Subsequently, at the indicated time points, cells were lysed in sterile water and serially diluted in PBS-Tween 80 0.05 %. Dilutions were plated on Middlebrook 7H10 complete agar medium (Difco Laboratories). Colonies were counted after 3 to 4 weeks to determine CFU.

### Mouse infections

Bacillary suspensions of the different strains were adjusted to 100 viable cells in 100 ml phosphate buffer saline (PBS). Each animal was anesthetized and intratracheally inoculated with *M. tuberculosis* strains (*Mtb*H37Rv, *Mtb*Δ*fasR* and *Mtb*Δ*fasR*-c*fasR*). Infected mice were kept in cages fitted with microisolators connected to negative pressure. Groups of 10 mice were infected with the different *M. tuberculosis* strains. The inoculum doses were determined by enumerating CFU recovered from the lungs of three mice per strain, 24 h post-infection. Seven mice per group were killed at 60 days after infection and lungs removed and homogenized. Four dilutions of each homogenate were spread onto triplicate plates of Middlebrook 7H10 complete agar medium. The number of colonies was counted after 3 to 4 weeks.

### Infection of THP-1 cells and indirect immunofluorescence (IFI)

*M. tuberculosis* strains were covalently stained with FITC (Sigma-Aldrich). Briefly, 1× 10^9^ bacteria were washed twice with 0.1 M PBS and suspended in 1 ml of the same solution. FITC was added to a final concentration of 5 μg/ml and incubated in the dark for 1 h at 37 °C. Bacteria were washed gently with PBS until unincorporated dye was eliminated, and then were used to infect THP-1 cells. THP-1 cells were maintained in complete full RPMI medium. The day before infection 2×10^5^ THP-1 cells per well were plated in a 24-well plate with coverslips in full RPMI medium plus 50 nM PMA to allow differentiation of the cells to macrophages. On the day of infection, cells were washed and infected with the different stained *M. tuberculosis* strains (*Mtb*H37Rv, *Mtb*Δ*fasR* and *Mtb*Δ*fasR*-c*fasR*) prepared in RPMI medium, at a MOI of 5. After 2 h of uptake, cells were washed and incubated with 10 μM Lysotracker Red DND-99 for 1 hour to allow the staining of acidic compartments, fixed with 4 % paraformaldehyde solution in PBS for 30 min and subjected to immunofluorescence analysis.

Cells were quenched by incubation with 50 mM NH4Cl solution in PBS for 30 min. and permeabilized with 0.05 % saponin/1 % BSA in PBS for 15 min, followed by overnight incubation with goat anti-LAMP-3 (Santa Cruz Biotechnology, Santa Cruz, CA) primary antibody diluted 1:50 in PBS. Secondary anti-goat antibody conjugated to Cy3 (Jackson Immuno Research Labs, Inc.) was diluted 1:600 in PBS. Cells were mounted with mounting medium (Dako, Denmark) and analysed by confocal microscopy using an SP5 AOBS confocal microscope (Leica Microsystems, Germany). Mycobacterial internalization was monitored by following FITC fluorescence (green). Lysotracker Red DND-99 or LAMP-3 association with mycobacterial phagosomes was analysed in at least 250 cells using Fiji software (U.S. National Institutes of Health, Bethesda, MD) as described previously (Vázquez et al., 2017). The experiments were performed in duplicates in three independent experiments. The statistical analysis was performed using two-tailed Student’s t-test.

## Results

### Construction and characterization of *M. tuberculosis fasR* mutant strain

To investigate the function of FasR in *M. tuberculosis*, and considering that *fasR* is essential for *M. smegmatis* viability, we first constructed an *M. tuberculosis* conditional knockdown strain in *fasR* (Figure 1A). Plasmids pFR33, providing a second copy of *fasR* under the control of the TET-PIP OFF system at the chromosomal L5 phage attachment site *attB* (Boldrin et al., 2010) and pGA80, were simultaneously transformed in *M. tuberculosis* (Table S1). Disruption of *fasR* was then obtained by two-step allelic exchange using pFR35, which deleted the gene at the native locus. Homologous recombination was confirmed by colony PCR and Southern blot analysis, and the resulting conditional knockdown strain was named *Mtb*Δ*fasR*-c*fasR* (Figure 1B). In this strain, expression of *fasR* was downregulated by addition of anhydrotetracycline (ATc) to the culture medium, thus leading to protein depletion. However, we were not able to detect any growth defect after addition of ATc, suggesting that *fasR* was not essential for *M. tuberculosis* survival. This result prompted us to try to obtain a *fasR* knockout (KO) mutant strain by replacing the *fasR* complementing plasmid with the empty vector pND255. Consistent with the ability of the conditional mutant to grow in the non-permissive condition, plenty of colonies were obtained in the presence of Hyg, and the replacement was successfully confirmed by colony PCR (Figure S1A). We selected two Hyg-resistant clones and confirmed the depletion of FasR by Western blot using total protein extracts (Figure S1B). The genotype of this strain was further analyzed by Southern Blot (Figure 1B), thus validating the genomic deletion of *fasR* and the generation of the *fasR* KO mutant strain, named *Mtb*Δ*fasR*.

**Figure 1.**
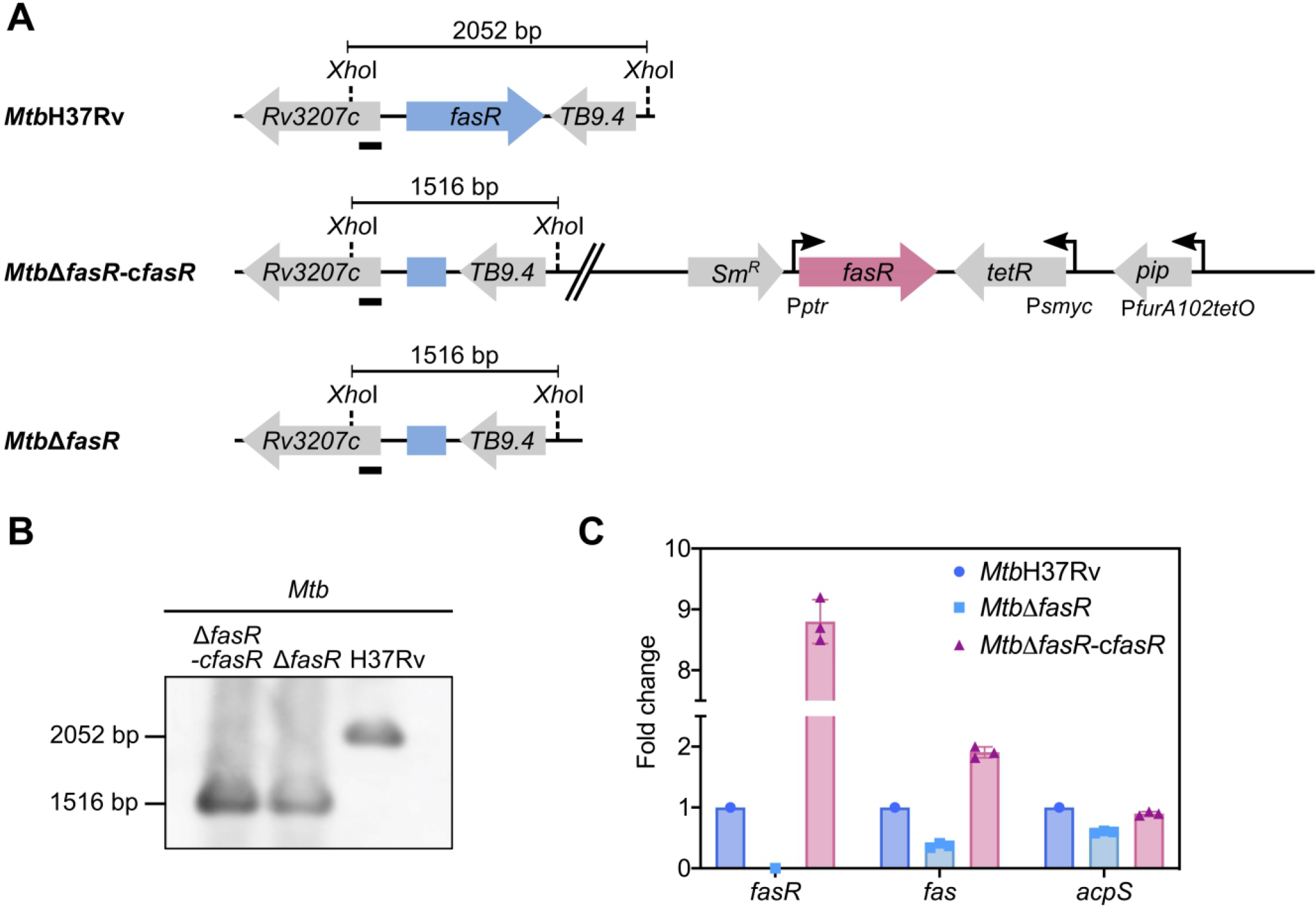
Construction and characterization of M. tuberculosis *fasR* mutant. (**A**) Schematic representation of the genetic organization of the chromosomal region of *fasR* in the wild type H37Rv strain (*Mtb*H37Rv), *fasR* conditional knockdown strain (*Mtb*Δ*fasR*-c*fasR*) and *fasR* KO strain (*Mtb*Δ*fasR*). **(B)** Southern Blot analysis of the different strains: chromosomal DNA was digested with *XhoI* and probed for hybridization with the probe indicated with a black bar, corresponding to the 5’ end of the *Rv3207* gene. **(C)** Relative expression levels of *fasR, fas* and *acpS* mRNAs measured by quantitative RT-PCR. Values were normalized to *sigA* and to the wild type strain (*Mtb*H37Rv). Samples for RNA extraction were collected at OD_600nm_ ~ 0.3.

We have previously demonstrated that FasR is a transcriptional activator of *fas* in *M. smegmatis* (Mondino et al., 2013). Thus, to validate the role of FasR in fatty acid biosynthesis in *M. tuberculosis*, we analyzed expression of the *fas-acpS* operon genes in *Mtb*Δ*fasR* and in the complemented strain *Mtb*Δ*fasR*-c*fasR*. The relative amounts of *fas* and *acpS* mRNAs were measured by quantitative RT-PCR after 8 days of growth in complete 7H9 medium and compared to those of *Mtb*H37Rv wild type strain. As shown in Figure 1C, transcription of *fas* and *acpS* was reduced by 70 % and 40 % respectively when cells were deprived of FasR, confirming that FasR is a transcriptional activator of the *fas-acpS* operon in *M. tuberculosis*.

Once the non-essentiality of *fasR* for *M. tuberculosis* survival was confirmed, we then hypothesized that deletion of this transcriptional regulator could be deleterious for bacterial growth under specific conditions. In particular, we analyzed the growth of *Mtb*Δ*fasR* mutant using different culture media. As shown in Figure 2A, growth of *Mtb*Δ*fasR* in complete 7H9 medium (7H9 + OADC) was similar to that of the wild type and the complemented strain (*Mtb*Δ*fasR*-c*fasR* grown without ATc), although the KO mutant reached stationary phase earlier and at lower OD_600_. Interestingly, if the medium was not supplemented with oleate and BSA (i.e. 7H9 without OADC), growth defects appeared to be more pronounced, suggesting that the presence of lipids in complete 7H9 medium could partially rescue the absence of FasR (Figure 2B). Moreover, *Mtb*Δ*fasR* mutant showed severe growth defects in Sauton minimal medium, reaching a maximal OD_600_ of 0.3 (Figure 2C). Altogether, these experiments showed that FasR is not essential for *M. tuberculosis* survival, although it is required for optimal bacterial growth, especially under lipid-limited conditions.

**Figure 2.**
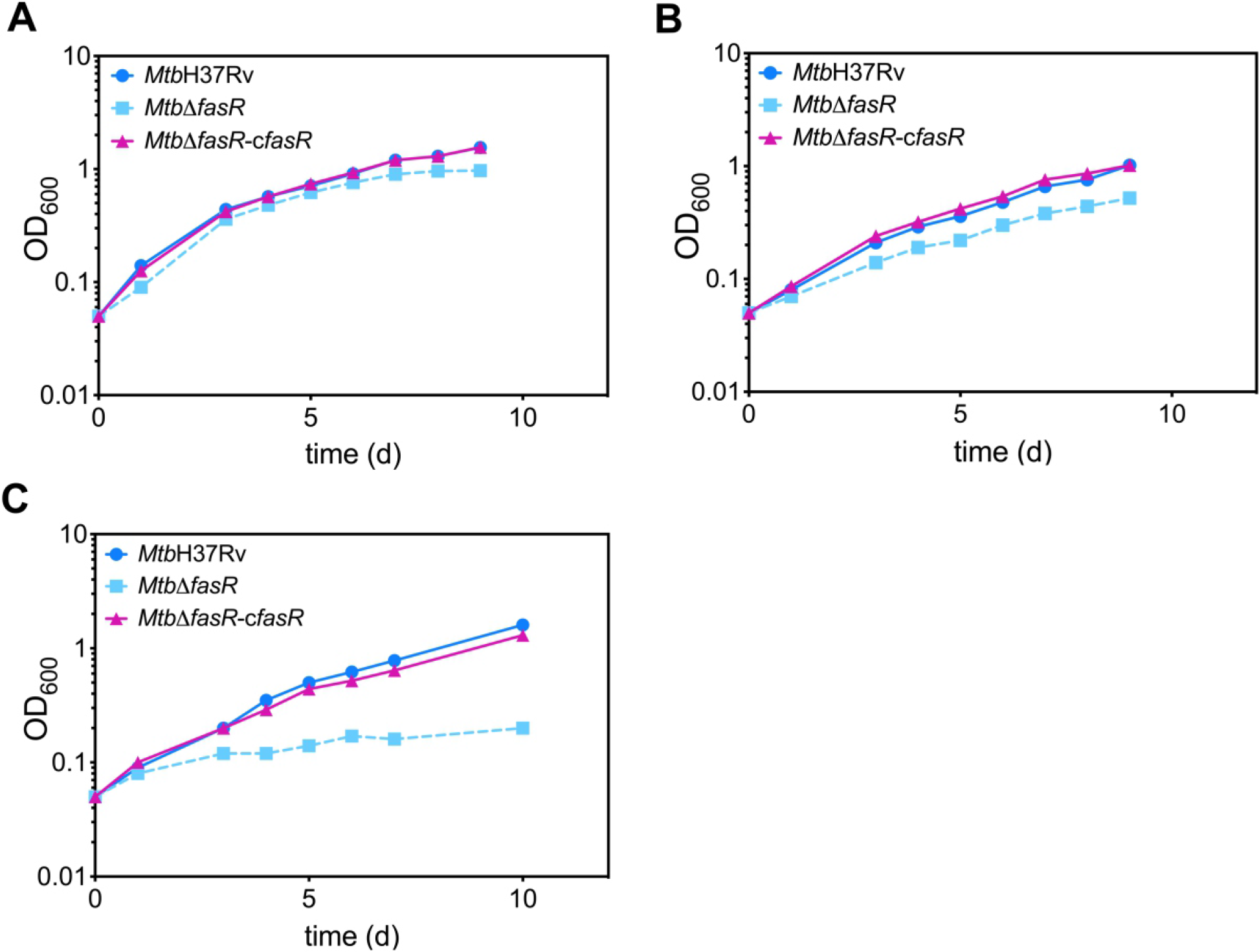
Growth of *M. tuberculosis* strains. Growth curves of the wild type strain (*Mtb*H37Rv), the *fasR Mtb*Δ*fasR*-c*fasR* strain grown without ATc (complemented strain) and *fasR* KO strain (*Mtb*Δ*fasR*) were obtained in 7H9 complete medium **(A)**, 7H9 without OADC medium **(B)** and Sauton medium **(C)**. Growth was followed by measuring OD_600 nm_. Results are representative of three independent experiments.

### *fasR* knockout strongly impacts composition of *M. tuberculosis* cell envelope

Previous characterization of *M. smegmatis fasR* (Mondino et al., 2013) and *fas* (Cabruja et al., 2017) conditional mutant strains revealed that reduced FAS I levels had a strong impact on fatty acid and phospholipid biosynthesis. Interestingly, MA remained being synthesized in the FAS I mutant for several hours after addition of ATc (non-permissive conditions), although with different abundance of the various species (Cabruja et al., 2017). Considering the relevance that lipids have in *M. tuberculosis* survival and virulence, the impact of the absence of FasR on the global lipid profile of *M. tuberculosis* was further studied. We performed comparative lipidomic analyses between *Mtb*H37Rv, *Mtb*Δ*fasR* and the complemented strain. Briefly, we extracted cell-associated lipids as previously described (Cabruja et al., 2017) and analyzed the lipid content by high-performance liquid chromatography-mass spectrometry with data-dependent tandem MS (HPLC-MS/MS). As illustrated in Figure 3, lipid analysis in the negative mode indicated that the relative abundance (expressed as percentage of lipid MS signals) of phospholipids (PL) and MA in the wild type and complemented strains was approximately 75 and 25 %, respectively. These values changed to 50 % for both PL and MA in the *fasR* mutant strain, reflecting the strong impact that lower levels of *de novo* fatty acid biosynthesis had in PL and MA relative abundance. The relative composition of PL also showed a biased effect towards phosphatidylinositol (PI) and cardiolipins (CL), which were significantly reduced in the mutant strain (Figures 3 and S2). Furthermore, there was a strong reduction in the content of sulfolipids (Figures 3 and S2) and the acyl chain substitutions of this lipid were consistently shorter in the mutant strain (Figure S3). A relevant change observed in the qualitative composition of the PL was the significant increase in the double bonds of the acyl residues in the mutant strain, with those of the CL, phosphatidylglycerol (PG) and PI being most significant (Figure S4).

**Figure 3.**
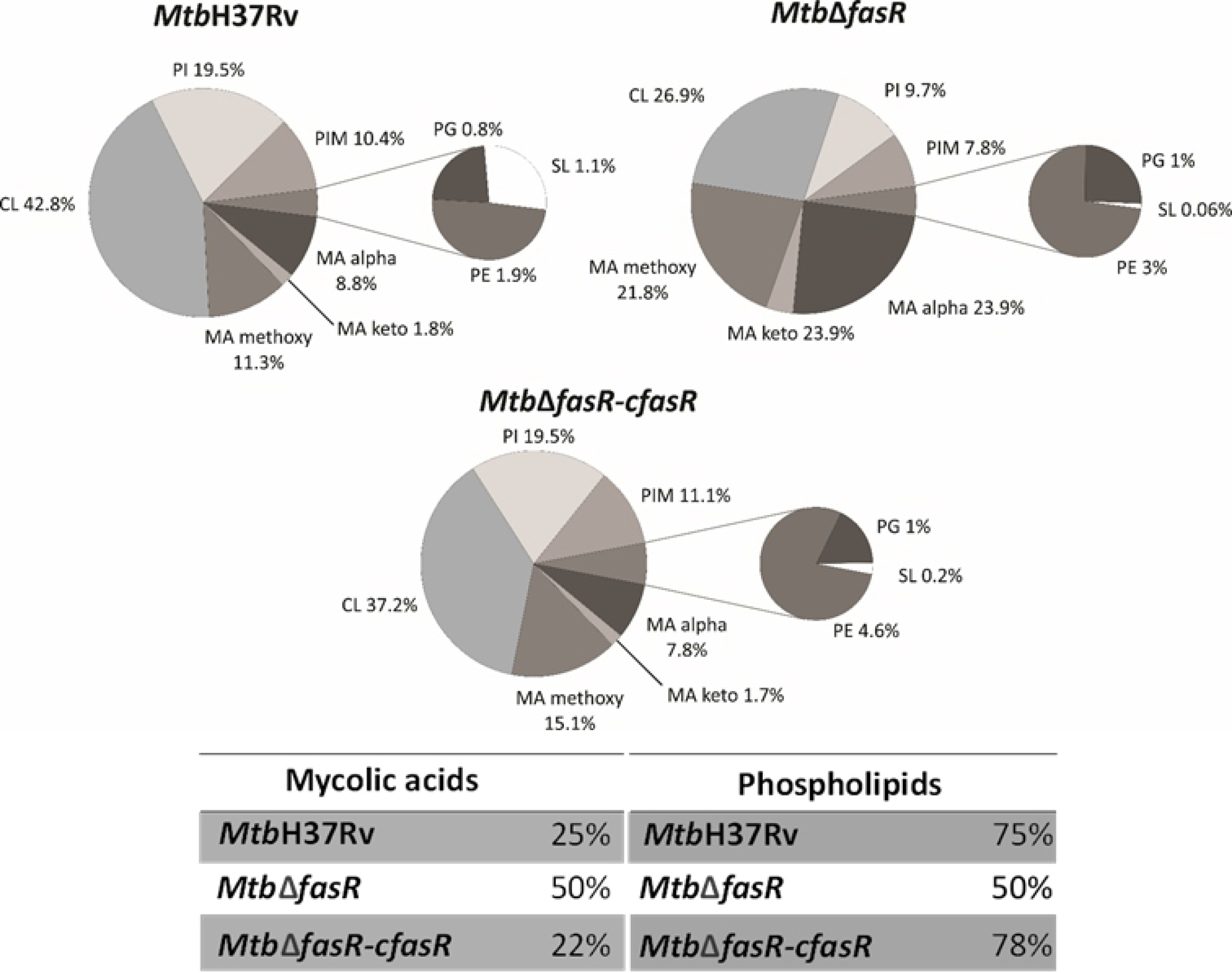
Lipid profile in negative ion mode. LC-MS analyses (negative mode) of the phospholipids and mycolic acids present in the wild type strain (*Mtb*H37Rv), the *fasR* complemented strain (*Mtb*Δ*fasR*-c*fasR*) and *fasR* KO strain (*Mtb*Δ*fasR*). The values indicated represent the relative abundance of the MS signals corresponding to the different lipid classes found in the samples. Results are the means of three independent experiments. CL, cardiolipin; PI, phosphatidylinositol; PIM, phosphatidylinositol mannosides; PG, phosphatidylglycerol; PE, phosphatidylethanolamine; SL, sulfolipids; MA, mycolic acids.

Additionally, results obtained in the positive mode analysis showed that *M. tuberculosis Mtb*Δ*fasR* mutant strain displayed a significant reduction in the relative content of PDIM (Figures 4 and S2), with acyl substitutions that are shorter in the mutant strain, compared with the wild type or the complemented strain (Figure S3). Most of the changes observed in the deletion mutant were reversed in the complemented strain *Mtb*Δ*fasR*-c*fasR*, indicating that *fasR* is needed to maintain the global lipid composition of *M. tuberculosis* cell envelope.

**Figure 4.**
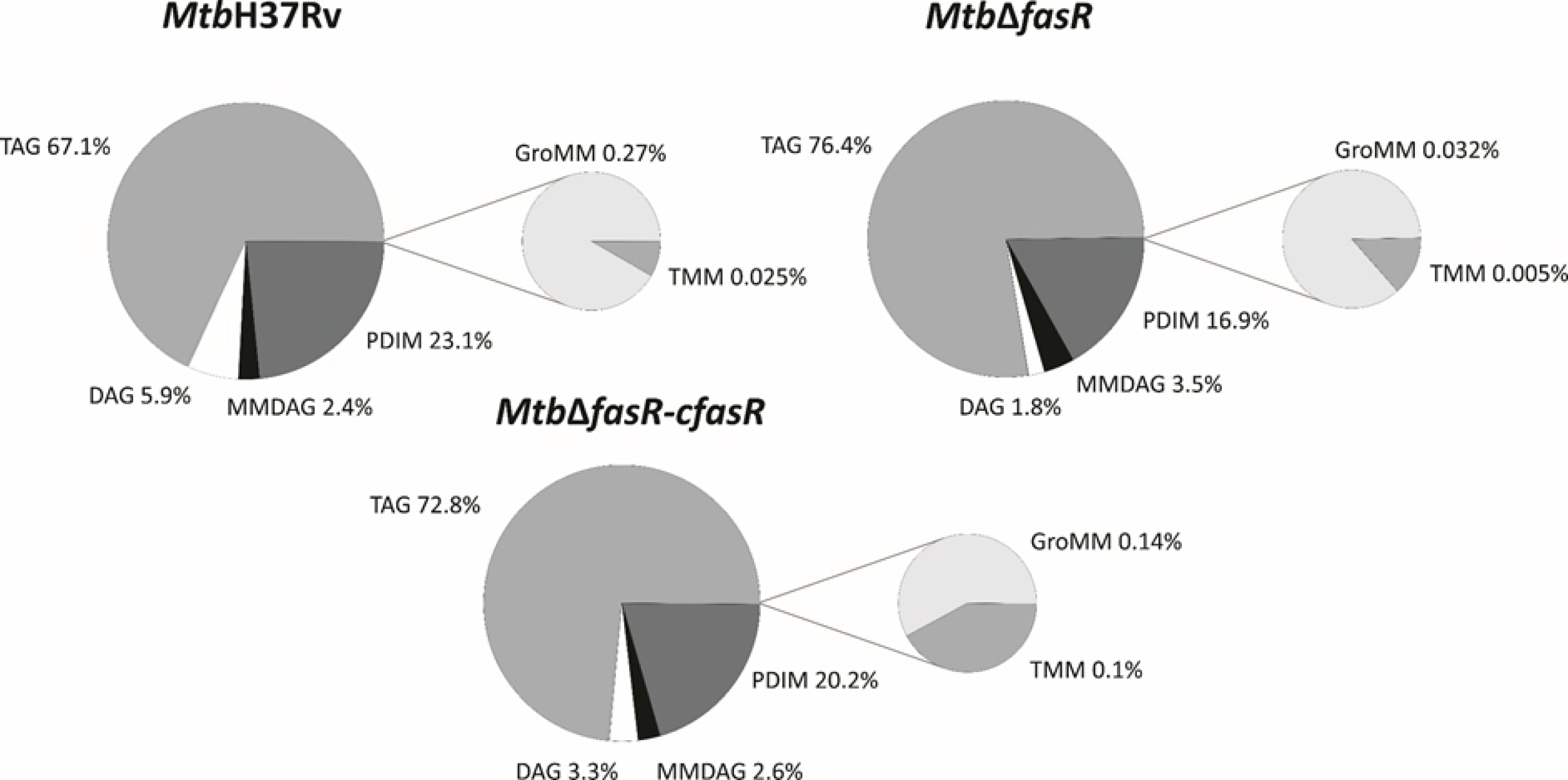
Lipid profile in positive ion mode. LC-MS analyses (positive mode) of lipids present in the wild type strain (*Mtb*H37Rv), the *fasR* complemented strain (*Mtb*Δ*fasR*-c*fasR*) and *fasR* KO strain (*Mtb*Δ*fasR*). The values indicated represent the relative abundance of the MS signals corresponding to the different lipid classes found in the samples. Results are the means of three independent experiments. TAG, triacylglycerol; DAG, diacylglycerol; MMDAG, monomycolyl DAG; PDIM, phthiocerol dimycocerosates; GroMM, glycerol monomycolate; TMM, trehalose monomycolate.

### FasR is required for *M. tuberculosis* replication in macrophages

To investigate whether FasR is required for the intracellular survival of *M. tuberculosis* in macrophages, we infected THP-1-derived macrophages with exponentially growing cultures of *Mtb*H37Rv, *Mtb*Δ*fasR* and the complemented strain *Mtb*Δ*fasR*-c*fasR*. To quantify bacterial uptake and intracellular replication, we determined the numbers of intracellular viable bacteria at different time points post-infection. As shown in Figure 5A, infection with similar numbers of bacteria of each strain resulted in recovery of similar numbers of CFU for all of the strains at 24 and 48 h post-infection. However, 120 and 144 h after the infection, intracellular bacterial replication of *Mtb*Δ*fasR* strain was significantly reduced compared to that of the wild type strain. The wild type phenotype was partially restored in the *Mtb*Δ*fasR*-c*fasR* complemented strain at day 5 (120 h) and completely restored at day 6 (144 h), indicating that the intracellular growth defect was due to the deletion of *fasR* and that this regulator is necessary for optimal intracellular multiplication of *M. tuberculosis* in macrophages.

**Figure 5.**
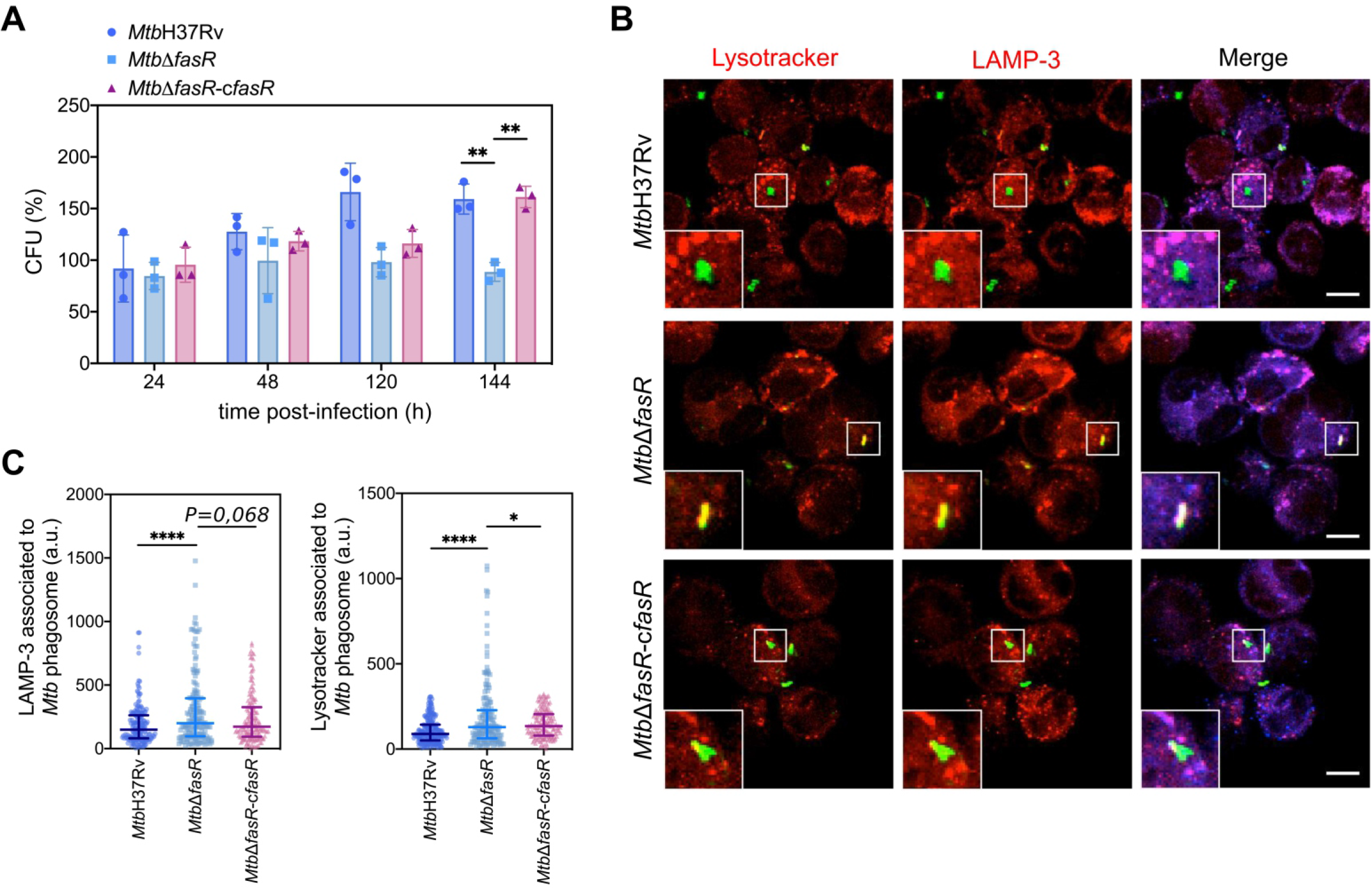
FasR is required for the intracellular survival of *M. tuberculosis*. **(A)** THP-1-derived macrophages were infected with wild type *M. tuberculosis* H37Rv (*Mtb*H37Rv), *fasR* mutant (*Mtb*Δ*fasR*) and the complemented *fasR* strain (*Mtb*Δ*fasR*-c*fasR*). At different times post-infection (24, 48, 120 and 144 hours) intracellular viable bacteria were counted. Bars represent means ± standard deviation of three independent experiments. Data were analyzed using two-way ANOVA with Tukey’s multiple comparisons test (** P≤0,01) **(B)** Representative confocal images of THP-1 macrophages infected with the different FITC-labelled strains. Acidic compartments were stained with Lysotracker Red DND-99 and LAMP-3 was detected by indirect immunofluorescence using a specific antibody. Scale bars: 10 μm. Merge indicates the location of the different strains in the phagolysosome. **(C)** Fluorescence intensity quantification of Lysotracker and LAMP-3 associated with the phagosome of *M. tuberculosis Mtb*H37Rv (circles), *Mtb*Δ*fasR* (squares) and the complemented *fasR* strain *Mtb*Δ*fasR*-c*fasR* (triangles). Data were expressed in arbitrary units (a.u.). Fiji software was used for quantification. Data represent medians with interquartile range of three independent experiments. Data were analyzed using Brown-Forsythe and Welch ANOVA tests with Games-Howell’s multiple comparisons test (*, P ≤ 0.05; ****, P ≤ 0.0001).

### FasR regulates phagosome maturation in macrophages

We have shown that the absence of FasR, and consequently reduced levels of *fas-acpS*, have a strong impact on the composition of *M. tuberculosis* cell envelope, and also on bacterial multiplication within macrophages. We hypothesized that the altered bacterial cell envelope of the *Mtb*Δ*fasR* mutant may affect *M. tuberculosis*-driven phagosome maturation during infection. To investigate this, we analyzed the acidification of *M. tuberculosis Mtb*Δ*fasR* phagosome during infection of human macrophages by staining with Lysotracker dye and measuring the phagosomal acquisition of LAMP-3 (late endosomal/lysosomal markers). Interestingly, while *Mtb*H37Rv prevented phagosome acidification, *Mtb*Δ*fasR* mutant strain resided in phagosomes that were strongly stained by Lysotracker dye and that acquired LAMP-3, suggesting that this strain cannot avoid phagosomal maturation. The observed mutant phenotype was rescued in the *Mtb*Δ*fasR*-c*fasR* complemented strain, indicating that the increased phagosome acidification was due to the absence of FasR (Figure 5B,C).

### Disruption of lipid homeostasis attenuates mycobacterial virulence in mice

To characterize the role of FasR in an experimental model of infection, three groups of BALB/c mice (10 per group) were infected intratracheally with 10^2^ CFU of *Mtb*H37Rv, *Mtb*Δ*fasR* and *Mtb*Δ*fasR*-c*fasR* strains. Mice were killed 60 days post-infection and bacterial loads in lungs were determined. CFU counts were significantly reduced for the *Mtb*Δ*fasR* strain compared to *Mtb*H37Rv, while the wild type phenotype was restored in the *Mtb*Δ*fasR*-c*fasR* strain (Figure 6), indicating that FasR is essential for survival and multiplication of *M. tuberculosis* in mice.

**Figure 6.**
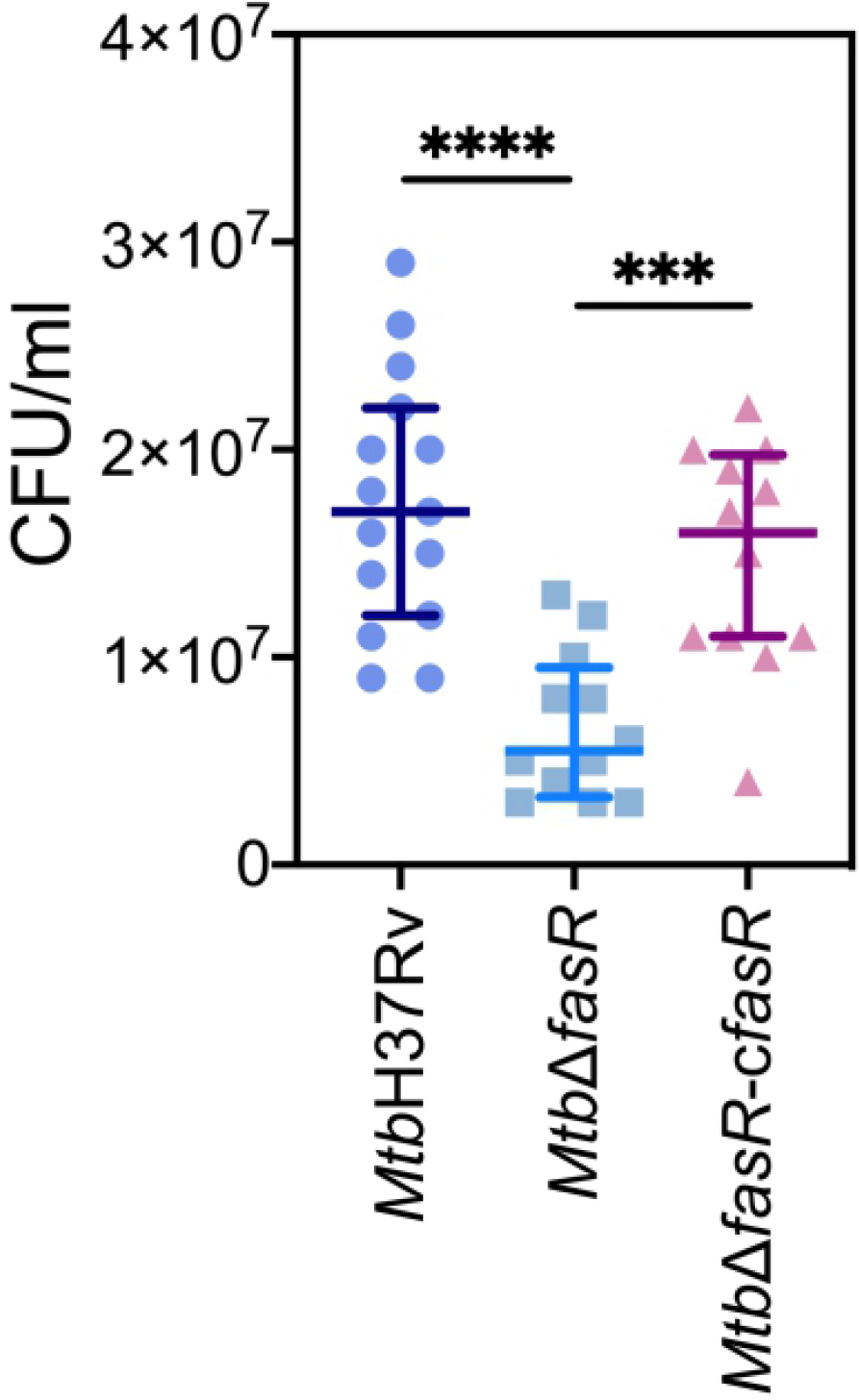
FasR is essential for the multiplication of mycobacteria in mouse lungs. Three groups of BALB/c mice (10 per group) were infected intratracheally with 10^2^ CFU of wild type *M. tuberculosis* H37Rv (*Mtb*H37Rv), *fasR* mutant (*Mtb*Δ*fasR*) and the complemented *fasR* strain (*Mtb*Δ*fasR*-c*fasR*). Bacterial loads in lungs were determined 60 days post-infection. Data represent medians with interquartile range. Data were analyzed using Brown-Forsythe and Welch ANOVA tests with Dunnett’s T3 multiple comparisons test (***, P ≤0.001; ****, P ≤ 0.0001).

## Discussion

In a previous investigation we identified FasR, a TetR-like transcriptional regulator that specifically binds to the *fas* promoter region to activate its expression, and found that FasR was essential for *in vitro* growth of *M. smegmatis* (Mondino et al., 2013). However, its role in *M. tuberculosis* viability was controversial. Using Himar 1 transposon mutagenesis it was initially proposed that FasR was essential for *M. tuberculosis* survival (Sassetti et al., 2003; Griffin et al., 2011). However, more recently, De Jesus *et al* found that FasR was dispensable for growth, by assessing the composition of a highly saturated library of transposon mutants *of M. tuberculosis* by deep sequencing (DeJesus et al., 2017). Moreover, the relevance of FasR for mycobacterial survival in intracellular environments or animal models had not been investigated. In this work, by constructing a deletion *fasR* mutant and a complemented strain, we unequivocally established that FasR is not essential for *M. tuberculosis* viability, although it is required for optimal growth of *M. tuberculosis in vitro* and for virulence in macrophages and mice model of infection.

We had shown that FasR binds to three inverted repeats present in the promoter region of the *fas-acpS* operon thereby activating fatty acid biosynthesis in *M. smegmatis* (Mondino et al., 2013). Furthermore, we also demonstrated, *in vitro* and *in vivo*, that FasR binding to its target promoter is regulated by long-chain acyl-CoAs, the main products of FAS I (Mondino et al., 2013). The *fas* and *acpS* genes form a bicistronic operon, coding for the FAS I synthase and the 4-phosphopantetheinyl transferase, respectively; the latter being an essential enzyme to produce functional acyl carrier protein (holo ACP), central to all fatty-acid biosynthesis systems. Transcriptional studies carried out in this work showed that expression of the *fas-acpS* operon genes in *M. tuberculosis* Δ*fasR* was reduced by 70 % and 40 %, respectively (Figure 1C). This result confirmed that FasR is a transcriptional activator of the *fas-acpS* operon in *M. tuberculosis*. The activator nature of FasR highlights a remarkable difference with most transcriptional regulators of fatty acid biosynthesis, which are repressor proteins (Zhu et al., 2009). Moreover, FasR is a member of the TetR family of regulators, and the vast majority of these proteins are transcriptional repressors, with very few acting as activators (Cuthbertson and Nodwell, 2013).

Since fatty acids are major constituents not only of phospholipids but also of several complex lipids, inactivation of FasR was expected to modify synthesis of several components of the cell envelope essential for mycobacterial viability and/or virulence. Indeed, lipidomic analysis of *Mtb*Δ*fasR* and comparison with wild type and complemented strains proved this hypothesis. As expected, reduced levels of fatty acid biosynthesis in *Mtb*Δ*fasR* had a strong impact on relative abundance of phospholipids and their relative and qualitative composition (Figures 3 and S2). Phosphatidylinositol (PI) and cardiolipins (CL) were significantly reduced in the mutant strain, while the acyl chains present in and these phospholipids exhibited a significant increase in their double bond content (Figure S4). Mycobacterial phospholipids have been implicated in immunomodulation. For example, CL is processed into lysocardiolipin by the lysosomal phospholipase A2 during infection, probably generating active lipid derivatives that play important roles during infection or persistence within the host (Fischer et al., 2001). The *Mtb*Δ*fasR* mutant strain also displayed a strong reduction in the content of sulfolipids (Figures 4 and S2) and PDIM (Figures 4 and S2). Interestingly, the acyl chain substitutions of both lipids were consistently shorter compared to the wild type *Mtb*H37Rv strain (Figure S3). On this regard, a *M. tuberculosis* mutant strain deficient in sulfolipids and PDIM has been shown to be attenuated in macrophages and in mice models of infection (Passemar et al., 2014). Moreover, during infection, genes involved in the biosynthesis of sulfolipids and PDIM are upregulated (Graham and Clark-Curtiss, 1999; Rodríguez et al., 2013) and the mass of these lipids increases (Yang et al., 2009; Griffin et al., 2012). Altogether, our results showed that *M. tuberculosis fasR* mutant exhibits an extensive rearrangement of the lipid components of the cell envelope, that are reversed in the complemented strain *Mtb*Δ*fasR*-c*fasR*, confirming that FasR is needed to maintain the lipid homeostasis of the *M. tuberculosis* cell envelope. As the mycobacterial cell envelope is the interface with the host, FasR modulation of its structure and consequent responsiveness to immune factors might be important in determining the state of an infection.

Consistent with this hypothesis, we have shown that FasR is necessary for optimal intracellular multiplication of *M. tuberculosis* in macrophages, therefore confirming the role of cell wall components in the ability of the pathogen to survive and multiply intracellularly (Garcia-Vilanova et al., 2019). Persistence of the tubercle bacillus inside cells relies upon inhibition of phagosome maturation and acidification (Rohde et al., 2007). In this context, several cell wall lipids have been shown to modulate host endocytic pathways. This is the case for Man-Lam, TDM, PIM and PDIM, which, besides their role in phagocytosis, also mediate intracellular trafficking and vacuole maturation arrest induced by *M. tuberculosis* (Passemar et al., 2014). More recently, PDIM and SL were shown to be involved in controlling autophagy-related pathways in human macrophages (Bah et al., 2020). Considering the changes observed in the PDIM content of the *Mtb*Δ*fasR* mutant, we analyzed the acidification of phagosome during infection of human macrophages. Interestingly, while the wild type *Mtb*H37Rv and the complemented strains prevented phagosome acidification, the *Mtb*Δ*fasR* mutant was impaired in avoiding phagosomal maturation, clearly indicating that increased phagosome acidification was due to absence of FasR (Figure 5). These results suggested that although FasR is not essential for *in vitro* growth, regulation of lipid biosynthesis, mediated by FasR, is critical for macrophage infection. Moreover, using a mouse model of infection we demonstrated the essentiality of FasR for *M. tuberculosis* virulence *in vivo* (Figure 6), most probably as a consequence of the loss of various mycobacterial lipids that contribute to pathogenicity.

Co-evolution with the human immune system prompted *M. tuberculosis* to generate exquisite mechanisms of immunomodulation and immune evasion, many of which depend on its cell envelope. In fact, comparison of cell envelope components between *M. tuberculosis* and ancestral species revealed a restructuring of the mycomembrane glycolipid components, including mycolic acids and free lipids of the outer leaflet (Dulberger et al., 2020). The observed phenotype of the FasR mutant is probably a consequence of additive effects, as several components of the cell envelope were altered. *M. tuberculosis* interferes with innate immune signalling pathways, for example by masking pathogen-associated molecular patterns with PDIM (Cambier et al., 2014) and inhibiting toll-like receptor 2 with sulfoglycolipids (Blanc et al., 2017). PDIM is also crucial for other steps of infection: it contributes to cell envelope properties, such as the permeability barrier (Camacho et al., 2001), improves the invasiveness of *M. tuberculosis* and its capacity to arrest phagosome maturation, possibly by inducing changes in the plasma membranes of human macrophages during infection (Astarie-Dequeker et al., 2009), and directly protects against reactive nitrogen species (Rousseau et al., 2004). Therefore, reduced content of these molecules (and/or changes in their chemical composition) may contribute to sensitivity to the bactericidal activities of phagocytes.

Altogether, our data indicate an important role for FasR in *M. tuberculosis* pathogenesis and encourage further investigations into the contribution of lipid regulation to virulence. It is well known that *M. tuberculosis* is able to modulate the expression of its cell wall components in response to environmental changes, however, little is known about their reorganization during infection, and to a lesser extent about the regulatory mechanisms involved in this process (Dulberger et al., 2020). The dynamism of the bacterial cell envelope should be a tightly regulated phenomenon, ensuring *M. tuberculosis* survival during infection. Thus, specific regulatory components involved in this process, as we have demonstrated for FasR, would constitute interesting targets for the development of new treatment strategies.

Along this line of thinking, we recently solved the crystal structures of *M. tuberculosis* FasR in complex with DNA and the acyl effector ligands, which provided insights into the molecular sensory and transmission mechanisms of this regulatory protein (Lara et al., 2020). This knowledge could lead now to perform structure-guided drug discovery strategies to identify anti-mycobacterial molecules with novel mechanisms of action.

## Funding/Acknowledgements

This research was supported by Agencia Nacional de Promoción de la Investigatión, el Desarrollo Tecnológico y la Innovación (ANPCyT) grants 2015-0796 and 2018-02539 to GG and 2015-2022 to HG and by National Institutes of Health grant 1R01AI095183-01 to HG. The funders had no role in study design, data collection and analysis, decision to publish, or preparation of the manuscript.

