## Supplemental Information for "FasR regulates fatty acid biosynthesis and is essential for virulence of *Mycobacterium tuberculosis*"

**Supporting information**

**Table S1. Plasmids used in this study**

| Name | Description and features | References |
|---|---|---|
| pFR3 | pET28a(+) with *fasR* His tag fusion gene, under the control of T7 promoter, $Kan^R$ | (Mondino et al., 2013) |
| pJG1100 | Suicide vector for mutant construction, $Hyg^R$, $Kan^R$, *sacB* | (Gomez and Bishai, 2000) |
| pFR35 | Suicide vector for *Mtb*Δ*fasR* mutant construction derived from pJG1100, $Hyg^R$, $Kan^R$, *sacB* | This study |
| pGA44 | Integrative vector at L5 *attB* site, carrying the TetR/Pip OFF expression system and Ptr promoter, $Str^R$/$Spect^R$<br>The *int* gene was removed in order to lock this plasmid at the integration site. Integration requires plasmid pGA80 | (Kolly et al., 2014) |
| pGA80 | pMV261-derived vector, carrying the L5 *int* gene for expression in *trans*, lacking the mycobacterial origin of replication (Δ*oriM*), $Kan^R$ | (Kolly et al., 2014) |
| pFR33 | pGA44-derived vector carrying *fasR* under the control of the Ptr promoter, $Str^R$/$Spect^R$ | This study |
| pND255 | L5-integrative vector for mutant construction, $Hyg^R$ | (Kolly et al., 2014) |

$Hyg^R$, hygromicin resistance; $Kan^R$, kanamycin resistance; $Str^R$/$Spect^R$, streptomycin/spectinomycin resistance

**Table S2. Strains used in this study**

| Name | Description or Genotype | References |
|---|---|---|
| DH5α | *E. coli K12 F- ΔlacU169 (Φ80lacZΔM15) endA1 recA1 hsdR17 deoR supE44 thi-1λ- gyrA96 relA1* | (Hanahan, 1983) |
| *Mtb*H37Rv | *M. tuberculosis* wild type strain | (Cole et al., 1998) |
| *Mtb*Δ*fasR* | *Mtb*H37Rv background. Deletion of *fasR* | This study |
| *Mtb*Δ*fasR-cfasR* | *fasR* knockout complemented with *fasR* expressed *in trans* | This study |

**Table S3. Oligonucleotides used in this study**

| Name | Sequence (5’-3’) |
|---|---|
| 3208_1up | CTCGAGCCGGAATTCGTAGCC |
| 3208_2down | CATATGGAGCGCTGCCGGACGATCTG |
| 3208_3up | CATATGAAGCCGATTTCCAAGTCCGAC |

| | |
|---|---|
| 3208_4down | ACTAGTTGCCACCGTGGTGACCGTCTC |
| Southern_1fasRTB | GAACACTTTGACGGTGCCTTG |
| Southern_2fasRTB | GGAATGACATTACTACCTGCG |
| L-*sigA* | AAACAGATCGGCAAGGTAGC |
| R-*sigA* | CTGGATCAGGTCGAGAAACG |
| L-*fasR* | TGAGCACGACTACCGACAAC |
| R-*fasR* | CTCGAAGATCAGCCGGTAAC |
| L-*fas* | GACATCCTGACCCGACTGAC |
| R-*fas* | TGAACTTCGTCGAGAGCTTG |
| fasRF | ATGAGCGATCTCGCCAAGACAG |
| fasRR | CTACGAGCGGGTAAGCGGGACG |

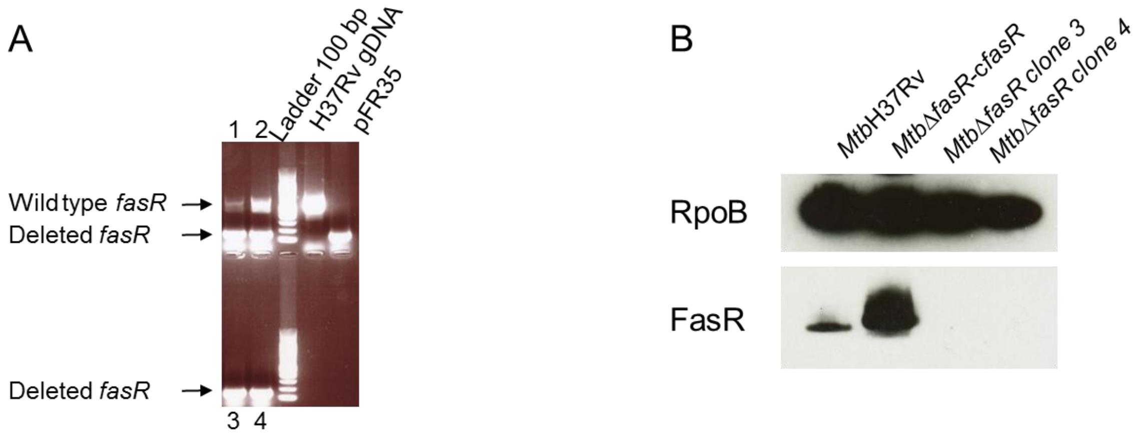


**Figure S1. Analysis of the knockout *fasR* mutant (A)** Colony PCR of *M. tuberculosis* transformants grown on Str, carrying the complementing *fasR* gene (clones 1 and 2) and of transformants grown on Hyg, carrying the empty pND255 vector (clones 3 and 4). Primers used for the amplification of *fasR* are listed in Table S3. The expected size of the PCR products is 687 bp for the *fasR* wild type allele and 150 bp for the mutant *fasR* allele. Two fragments were amplified from the Str-resistant clones, corresponding to the complementing *fasR* gene from the integrated plasmid and the deleted *fasR* gene at the *fasR* locus; on the other hand, only one fragment was amplified from the Hyg resistant clones (the partially deleted *fasR* gene) **(B)** Western Blot analysis of crude protein extracts prepared from cultures of wild type H37Rv strain (*Mtb*H37Rv), *fasR* conditional knockdown strain (*Mtb*Δ*fasR*-c*fasR*) and *fasR* KO strain (*Mtb*Δ*fasR* clones 3 and 4) grown at $OD_{600nm}$ ~ 0,3. Detection was performed using monoclonal anti-$FasR_{MT}$ antibodies (lower panel) and anti-RpoB as loading control (upper panel).

| Lipid profile in negative ion mode: signal % | | | | |
|---|---|---|---|---|
| | *MtbH*37Rv | *MtbΔfasR* | *MtbΔfasR-cfasR* | *p-value** |
| CL | 36.37 ± 2.43 | 20.87 ± 3.34 | 31.52 ± 0.78 | 0.0234 |
| PE | 1.54 ± 1.43 | 2.31 ± 1.25 | 3.92 ± 1.23 | 0.624 |
| PG | 0.74 ± 0.11 | 0.82 ± 0.07 | 0.86 ± 0.25 | 0.4725 |
| PI | 16.71 ± 2.26 | 7.49 ± 0.09 | 16.45 ± 3.35 | 0.0234 |
| PIMs | 8.97 ± 2.43 | 6.01 ± 1.06 | 9.35 ± 2.74 | 0.1851429 |
| MA alpha | 7.54 ± 1.43 | 18.44 ± 0.61 | 6.7 ± 2.2 | 0.018 |
| MA keto | 1.54 ± 0.22 | 2.98 ± 0.42 | 1.5 ± 0.62 | 0.0234 |
| MA methoxy | 9.66 ± 2.3 | 16.78 ± 0.22 | 13 ± 8.58 | 0.03825 |
| SL | 0.89 ± 0.71 | 0.04 ± 0.03 | 0.16 ± 0.04 | 0.0234 |

* T-test *MtbΔfasR vs MtbH*37Rv. Benjamini-Hochberg adjusted *p-value*.

| Lipid profile in positive ion mode: signal % | | | | |
|---|---|---|---|---|
| | *MtbH*37Rv | *MtbΔfasR* | *MtbΔfasR-cfasR* | *p-value** |
| DAG | 5.91 ± 1.37 | 1.84 ± 1.28 | 3.29 ± 0.19 | 0.0538 |
| TAGs | 67.15 ± 2.5 | 76.39 ± 0.61 | 72.78 ± 4.64 | 0.0246 |
| MMDAG | 2.43 ± 0.19 | 3.54 ± 0.25 | 2.57 ± 0.12 | 0.0246 |
| GroMM | 0.27 ± 0.13 | 0.03 ± 0.01 | 0.14 ± 0.047 | 0.0898 |
| PDIM | 23.1 ± 1.56 | 16.94 ± 0.24 | 20.22 ± 4.39 | 0.0246 |
| TMM | 0.02 ± 0.004 | 0.005 ± 0.001 | 0.1 ± 0.01 | 0.0246 |

* T-test *MtbΔfasR vs MtbH*37Rv. Benjamini-Hochberg adjusted *p-value*.

**Figure S2. Relative abundance of lipid species.** The tables show the relative abundance as the total signal percentage of the different lipid classes. CL, cardiolipin; PE, phosphatidylethanolamine; PG, phosphatidylglycerol; PI, phosphatidylinositol; PIMs, phosphatidylinositol mannosides; SL, sulfoglycolipids; MA, mycolic acids; DAG, diacylglycerol; TAGs, triacylglycerol; MMDAG, monomycolyl-DAG; GroMM, glycerol monomycolate; PDIM, phthiocerol dimycocerosates; TMM, trehalose monomycolate. Results are the mean of three independent experiments ± SD (n = 3). p values were calculated with unpaired *t* test with Benjamini-Hochberg's correction between *Mtb*Δ*fasR* and *Mtb*H37Rv.

A)

| Lipid profile in negative ion mode: C – chain length | | | | |
|---|---|---|---|---|
| | *MtbH37Rv* | *MtbΔfasR* | *MtbΔfasR-cfasR* | *p-value** |
| CL | 68.71 ±0.02 | 68.65 ±0.14 | 68.95 ±0.05 | 0.017 |
| PE | 34.4 ±0.28 | 34.32 ±0.08 | 34.33 ±0.21 | 0.909 |
| PG | 34.2 ±0.11 | 34.17 ±0.17 | 34.3 ±0.03 | 0.219 |
| PI | 34.93 ±0.04 | 34.96 ±0.12 | 34.99 ±0.04 | 0.711 |
| MA alpha | 78.64 ±0.004 | 78.37 ±0.04 | 78.52 ±0.09 | 0.078 |
| MA keto | 84.64 ±0.03 | 83.65 ±0.28 | 84.3 ±0.08 | 0.016 |
| MA methoxy | 85.34 ±0.34 | 84.77 ±0.02 | 85.12 ±0.05 | 0.002 |
| SL | 151.04 ±0.27 | 144.01 ±1.61 | 148.84 ±0.72 | 0.013 |

| Lipid profile in positive ion mode: C – chain length | | | | |
|---|---|---|---|---|
| | *MtbH37Rv* | *MtbΔfasR* | *MtbΔfasR-cfasR* | *p-value** |
| DAG | 35.49 ±0.09 | 36.82 ±0.96 | 35.28 ±0.06 | 0.031 |
| GroMM | 47.61 ±0.15 | 46.67 ±0.9 | 46.49 ±0.93 | 0.018 |
| TAGs | 55.87 ±0.21 | 58.65 ±0.43 | 56.34 ±0.33 | 0.003 |
| TMM alpha | 82.16 ±0.13 | 81.22 ±0.13 | 80.89 ±0.4 | 0.237 |
| TMM keto | 87.07 ±0.13 | 86.07 ±0.65 | 86.69 ±0.15 | 0.106 |
| TMM methoxy | 87.51 ±0.12 | 86.39 ±0.31 | 87.32 ±0.47 | 0.052 |
| PDIM A | 93.19 ±0.26 | 91.17 ±0.21 | 92.65 ±0.03 | 0.0005 |

* T-test *MtbΔfasR vs MtbH*37Rv. Benjamini-Hochberg adjusted *p-value*.

B)

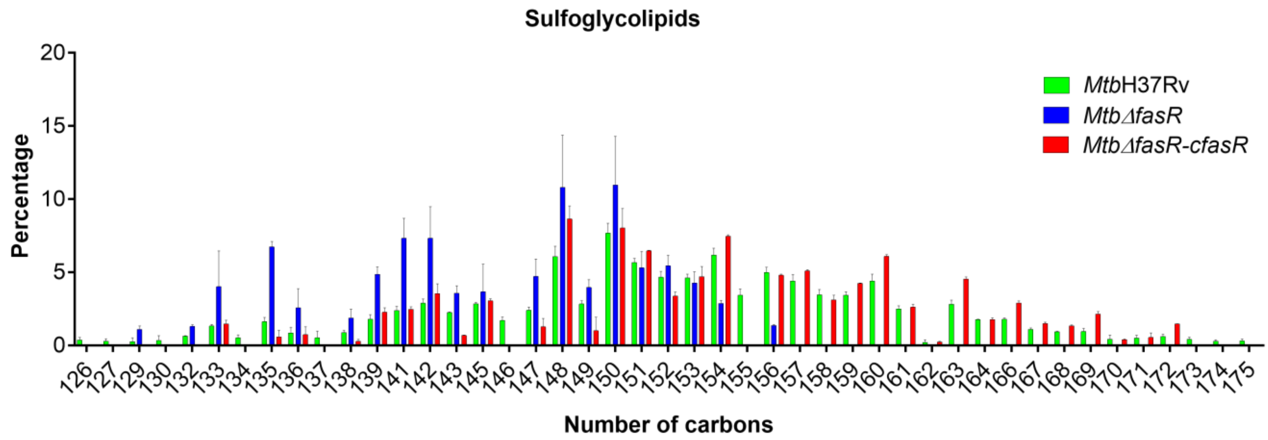


C)

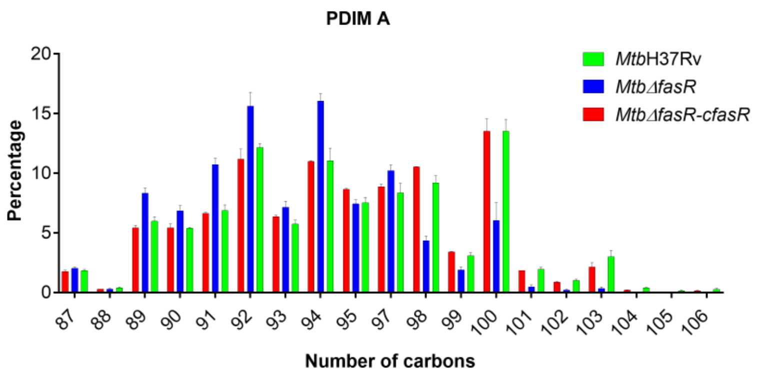

**Figure S3. Detailed analysis of lipid species. (A)** The tables show the weighted average of total number of carbons atoms per molecule within the population of mycolic acids (MA), phospholipids (PE, phosphatidylethanolamine; CL, cardiolipin; PG, phosphatidylglycerol; PI, phosphatidylinositol); sulfoglycolipids (SL); DAG, diacylglycerol; GroMM, glycerol monomycolate; TAGs, triacylglycerol; TMM trehalose monomycolate and phthiocerol dimycocerosates (PDIM A). The histograms show the distribution of the number of carbon atoms of SL **(B)** and PDIM **(C)** present in the wild type strain (*Mtb*H37Rv), the *fasR* complemented strain (*Mtb*Δ*fasR*-c*fasR*) and *fasR* mutant strain (*Mtb*Δ*fasR*). Results are the mean of three independent experiments ± SD (n = 3). p values were calculated with unpaired *t* test with Benjamini-Hochberg's correction between *Mtb*Δ*fasR* and *Mtb*H37Rv.

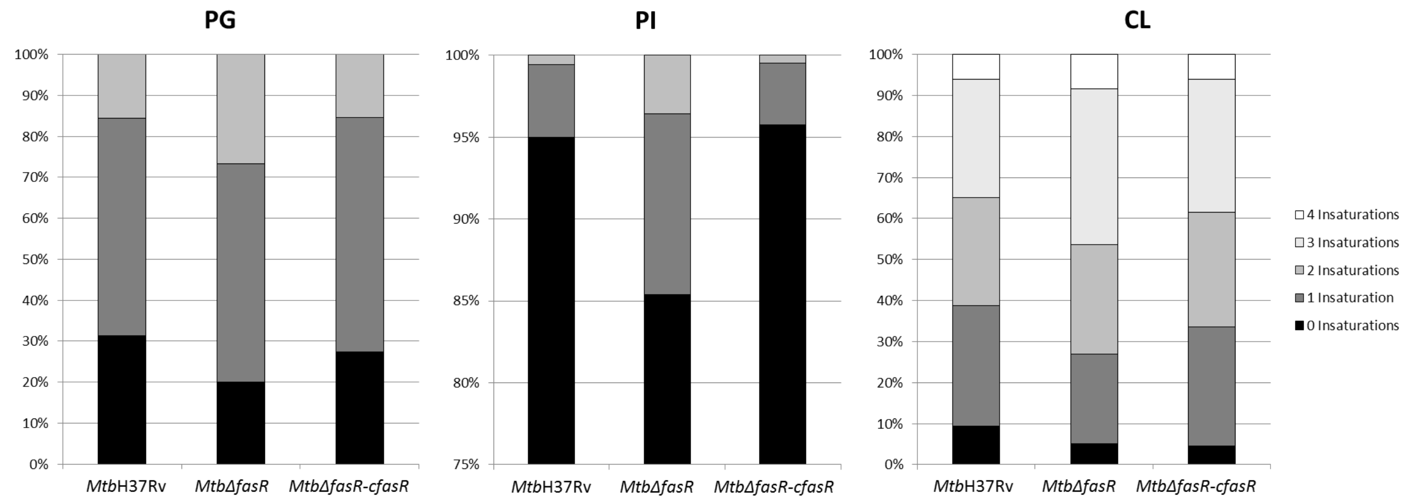


**Figure S4. Saturation level of the fatty acid substituents of the phospholipids.** Histograms represent the levels of unsaturation of the fatty acids constituents of cardiolipin (CL), phosphatidylglycerol (PG) and phosphatidylinositol (PI). Results are the means of three independent experiments.
